# Role of the H subunit C-terminal domain in the assembly of the vacuolar H+-ATPase

**DOI:** 10.1101/391656

**Authors:** Stuti Sharma, Rebecca A. Oot, Stephan Wilkens

**Affiliations:** Department of Biochemistry and Molecular Biology, SUNY Upstate Medical University 750 East Adams Street, Syracuse, New York 13210; Present address: Department of Cell Biology, Harvard Medical School, 250 Longwood Avenue, Boston, MA 02115

**Keywords:** Vacuolar HM^+^-ATPase, V_1_-ATPase, V_o_ proton channel subunit H, reversible disassembly: biolayer interferometry, native lipid nanodiscs

## Abstract

The vacuolar H^+^-ATPase (V-ATPase) is regulated by reversible disassembly into autoinhibited V_1_-ATPase and V_o_ proton channel sectors, a process that is poorly understood on the molecular level. V-ATPase is a rotary motor and recent structural analysis revealed that disassembled V_1_ and V_o_ are in different rotary states, a mismatch that is likely responsible for the inability to reconstitute holo V-ATPase from its functional sectors *in vitro*. Here, using the model organism *S. cerevisiae*, we show that a key impediment for binding of autoinhibited V_1_ to V_o_ is the conformation of the inhibitory C-terminus of subunit H (H_CT_). Using biolayer interferometry and biochemical analysis, we show that selective disruption of H_CT_’s binding site on V_1_ allows *in vitro* assembly of a structurally and functionally coupled V-ATPase complex. The resultant mutant V-ATPase, however, does not disassemble as readily as the wild type enzyme, highlighting the importance of H_CT_’s conformation in the mechanism of reversible disassembly. These findings pave the way for identifying molecules that allow for therapeutic modulation of aberrant V-ATPase activity in the disease state.

## Introduction

The vacuolar H^+^-ATPase (V-ATPase, V_1_V_o_-ATPase) is an ATP dependent proton pump found on the endomembrane system of all eukaryotic organisms. This multisubunit nano-motor functions in acidifying subcellular compartments, and in certain specialized tissues, the extracellular environment. Acidification by the V-ATPase is required for vital cellular processes such as pH and ion homeostasis, protein trafficking, autophagy, endocytosis, mTOR and Notch signaling as well as bone remodeling, urine acidification, hormone secretion and neurotransmitter release (1). While complete loss of V-ATPase function is embryonic lethal in mammals, aberrant activity has been associated with widespread human diseases such as renal tubular acidosis (2), osteoporosis (3), neurodegeneration (4), diabetes (5), male infertility (6), and cancer (7), thereby establishing the V-ATPase as a potential drug target (8, 9). However, because of its essential and ubiquitous nature, global non-selective inhibition of V-ATPase is not a viable option. Instead, there is a need for targeted modulation of the enzyme’s activity, a goal that requires a detailed understanding of V-ATPase’s catalytic and regulatory mechanisms.

V-ATPase is a large (~ 1 MDa) membrane protein complex that can be divided into a cytosolic V_1_-ATPase and a membrane integral V_o_ proton channel. In yeast, V_1_ and V_o_ are made of A_3_B_3_(C) DE_3_FG_3_H and *ac_8_c’c”def*, respectively (10). ATP hydrolysis takes place at three out of six A-B interfaces of the catalytic A3B3 hexamer. Located within the central cavity of the hexamer is subunit D, which together with F, provides the functional link between V_1_ and V_o_. V_o_ is constituted by subunit *a* that can be divided into cytosolic N-terminal and membrane-integral C-terminal domains (a_NT_ and a_CT_), the *c, c*’ and *c*” subunits (‘proteolipids’) that form a ring (*c*-ring), and subunit *d*. V_1_ and V_o_ are held together by three peripheral stalks, composed of heterodimers of subunits E and G (EG1-3) that connect the top of V_1_ to subunits C, H and *a*_NT_ (**Fig. 1*A***). V-ATPase is a rotary motor enzyme and employs a catalytic mechanism that is shared with the F-, A- and A-/V-type ATPases (11). In V-ATPase, ATP hydrolysis driven rotation of the central rotor (DF*d* and the *c*-ring) is coupled to proton translocation at the interface of a_CT_ and the *c*-ring. During catalysis, the three peripheral stalks, in conjunction with C, H and a_NT_, resist the rotary torque, thus keeping the A_3_B_3_ hexamer static against a_CT_ for efficient energy coupling.

**Fig. 1.**
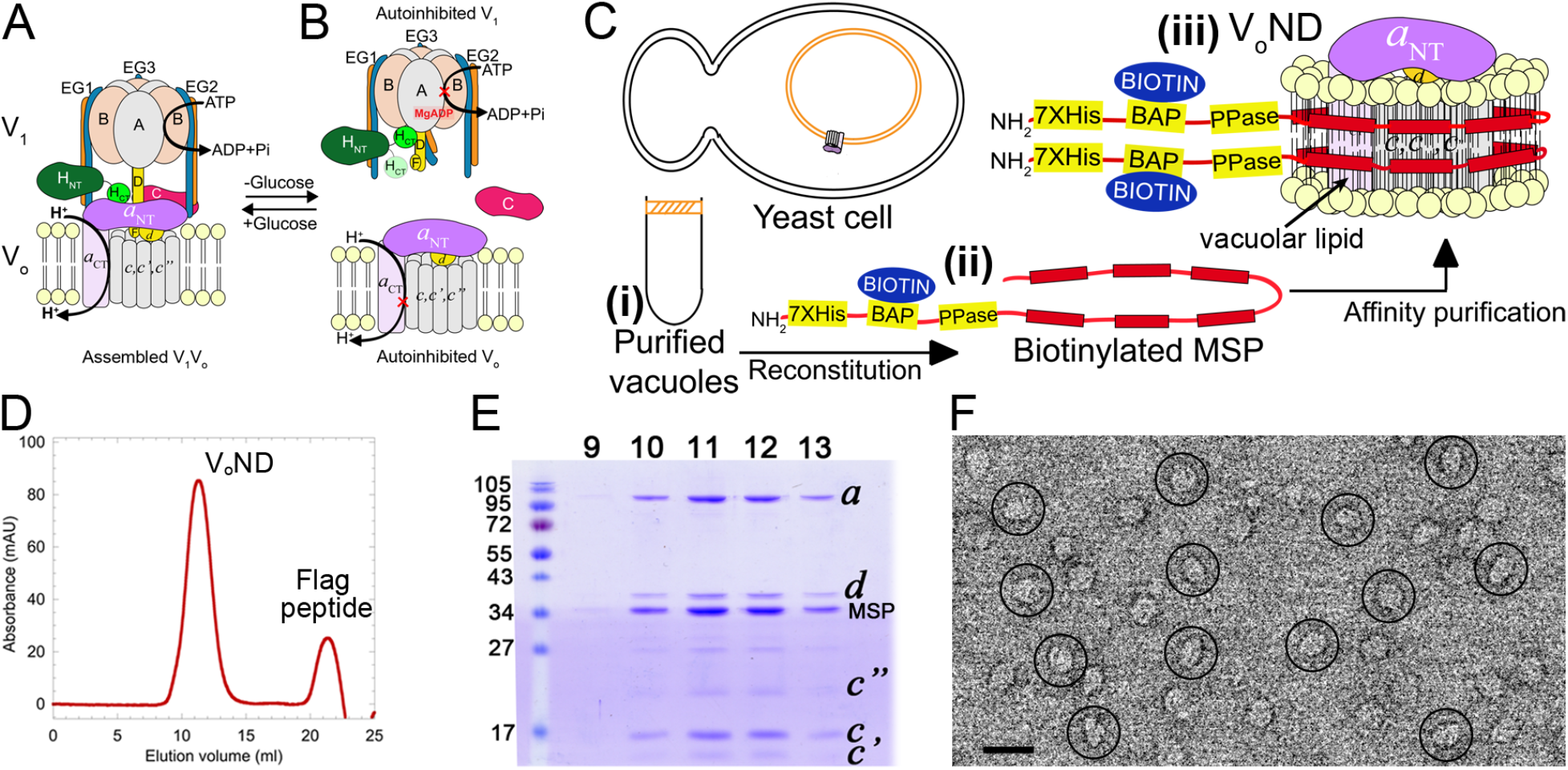
Purification and characterization of V_o_ND. A,B Schematic of V-ATPase regulation by reversible disassembly. C Purification strategy: Yeast vacuoles are isolated by flotation on a Ficoll gradient (i). Detergent solubilized vacuolar proteins are mixed with biotinylated MSP (ii) and reconstituted into lipid nanodiscs followed by α-FLAG affinity capture of V_o_ND (iii). D Size exclusion chromatography of V_o_ND. E Peak fractions were resolved using SDS-PAGE. F Negative stain EM of purified V_o_ND. Bar = 20 nm.

Unlike the related F- and A-ATPases, eukaryotic V-ATPase is regulated *in vivo* by a unique mechanism referred to as “reversible disassembly”, characterized by the detachment of V_1_ from V_o_ and the release of subunit C into the cytosol (12–14) (**Fig. 1*B***). The disassembly of V_1_ from V_o_, is accompanied by their “autoinhibition”, wherein free V_o_ no longer translocates protons (15, 16), and free V_1_ no longer hydrolyzes MgATP (17, 18). V-ATPase reversible disassembly was first described in yeast and insect, but is now emerging as an important regulatory mechanism in higher animals including human (19–21), and while there is some understanding of reversible disassembly at the cellular level (22), little is known about its molecular mechanism. Reversible disassembly requires the concerted breakage and reforming of subunit-subunit interactions, most of which involve the peripheral stator subunits E,G,H,C, and αNT. Using purified subunits and their domains, we found that while the head domain of C (C_head_) and the N-terminal domain of H (H_NT_) bind EG with nanomolar affinity, C’s foot domain (C_foot_), EG and H, all bind a_NT_ weakly (23, 24). It was therefore suggested that the V_1_-V_o_ interface is constituted by *multiple low-affinity* interactions that result in *high-avidity* binding. While the high-avidity allows for robust rotary catalysis, the low-affinity of the individual interactions permits efficient regulation of the enzyme by reversible disassembly (23, 25).

Recent structural studies revealed that in assembled V_1_V_o_, the C-terminal domain of H (H_CT_) is bound to a_NT_ (26), an interaction required for coupling V_1_’s ATPase to V_o_’s proton pumping activities (**Fig. 1*A***) (27). Upon disassembly of V_1_ from V_o_, H_CT_ undergoes a 150° rotation to wedge its inhibitory loop between the B subunit of an open catalytic site and the central stalk (DF) on V_1_ (**Supplementary Fig. S1*A-D***, red spheres). At the same time, *a*_NT_ moves from its peripheral position near Cfoot and EG that it occupies in V_1_V_o_ (28) (**Fig. 1*A***), towards a central position in autoinhibited V_o_ to bind subunit *d* (**Supplementary Fig. S1*E-G***) (16, 29–31). Moreover, a comparison of the cryoEM models of the three rotary states of holo V-ATPase (28) with the structures of autoinhibited V_1_ (32) and V_o_ (29, 31) showed that while V_1_ is halted in state 2, free V_o_ adopted state 3 (10) (**Supplementary Fig. S1**). From this observation, we hypothesized that the state mismatch together with the large conformational changes of H_CT_ and *a*NT that accompany enzyme dissociation explain why V_1_ does not readily bind free V_o_ under physiological conditions *in vitro* (10, 33), a safety-mechanism that likely evolved to prevent spontaneous reassembly *in vivo* when the disassembled state is required.

We recently introduced biolayer interferometry (BLI) of purified V-ATPase in biotinylated and endogenous vacuolar lipid containing nanodiscs to study enzyme dissociation (34). Here, we have expanded on this approach to study the interaction of V_1_ and V_o_. V_o_ sector was reconstituted into vacuolar lipid containing nanodiscs (V_o_ND) and immobilized on BLI sensors to screen V_1_ mutants for their ability to bind V_o_. We show that V_1_ in which wild type H is replaced with chimeric H composed of yeast H_NT_ and human H_CT_ (H_chim_; V_1_H_chim_) (32) was able to bind V_o_ND to form a structurally and functionally coupled V-ATPase (V_1_H_chim_V_o_ND). Characterization of V_1_H_chim_V_o_ND revealed biophysical and enzymatic properties similar to the previously characterized wild type V_1_V_o_ND (34). However, V_1_H_chim_V_o_ND was more resistant to ATP hydrolysis induced disassembly compared to wild type, highlighting the importance of H_CT_’s conformational switch in driving V-ATPase disassembly. The data also show that native vacuolar lipids are required for the generation of coupled MgATPase activity in reconstituted V_1_H_chim_V_o_ND. From the *in vitro* data presented here, we propose a molecular model for the reassembly of holo V-ATPase from autoinhibited V_1_ and V_o_ *in vivo*.

## Results

### Purification and characterization of native lipid nanodisc reconstituted V_o_ (V_o_ND) and V_1_ mutants

V_o_ was extracted from yeast vacuoles using the ‘reconstitution before purification’ strategy as described for V_1_V_o_ (34) (**Fig. 1*C*(i-iii)**). The resultant V_o_ND complex consisted of V_o_ embedded in endogenous vacuolar lipid containing nanodiscs encircled by biotinylated MSP (**Fig. 1*C*(iii)**). The purified complex was monodisperse and contained all V_o_ subunits plus MSP based on gel filtration and SDS-PAGE (**Fig. 1*D,E***). Examination of purified V_o_ND using negative stain electron microscopy showed single particles of V_o_ND with the typical size and appearance as described previously (30) (**Fig. 1*F***).

Four different V_1_ mutants were tested for their ability to bind V_o_ND and form coupled holo V-ATPase (**Fig. 2*A***): wild type V_1_, V_1_ purified from a yeast strain deleted for subunit H (V_1_ΔH), V_1_ΔH reconstituted with wild type H (V_1_H_wt_) and with chimeric H (H_chim_) consisting of yeast N-terminal and human C-terminal domains (32) (V_1_H_chim_). While wild type V_1_ has no measurable MgATPase activity (**Fig. 2*B***, dark green trace), V_1_ΔH had an initial specific activity of 15.7 ±1.7 U/mg, consistent with previous reports (32, 35) (**Fig. 2*B,C***, blue trace and bar). Although MgATPase activity is measured in an ATP regenerating system, the activity of V_1_ΔH decreases over time as MgADP gets trapped in a closed catalytic site, leading to the MgADP inhibited state (18, 24). V_1_H_wt_, which distinguishes itself from wild type V_1_ by having only ~0.4 instead of 1.3 mol/mol ADP in catalytic sites (32), exhibits an initial MgATPase activity of ~ 4.95 ± 0.55 U/mg before becoming MgADP inhibited (**Fig. 2*B,C*** light green). Reconstitution of V_1_ΔH with Hchim resulted in V_1_H_chim_ (**Fig. 2*D-F***), which showed an initial MgATPase activity of 8.2 ± 0.4 U/mg that, as for the other V_1_ mutants, declined over time due to MgADP inhibition (**Fig. 2*B,C*** pink).

**Fig. 2.**
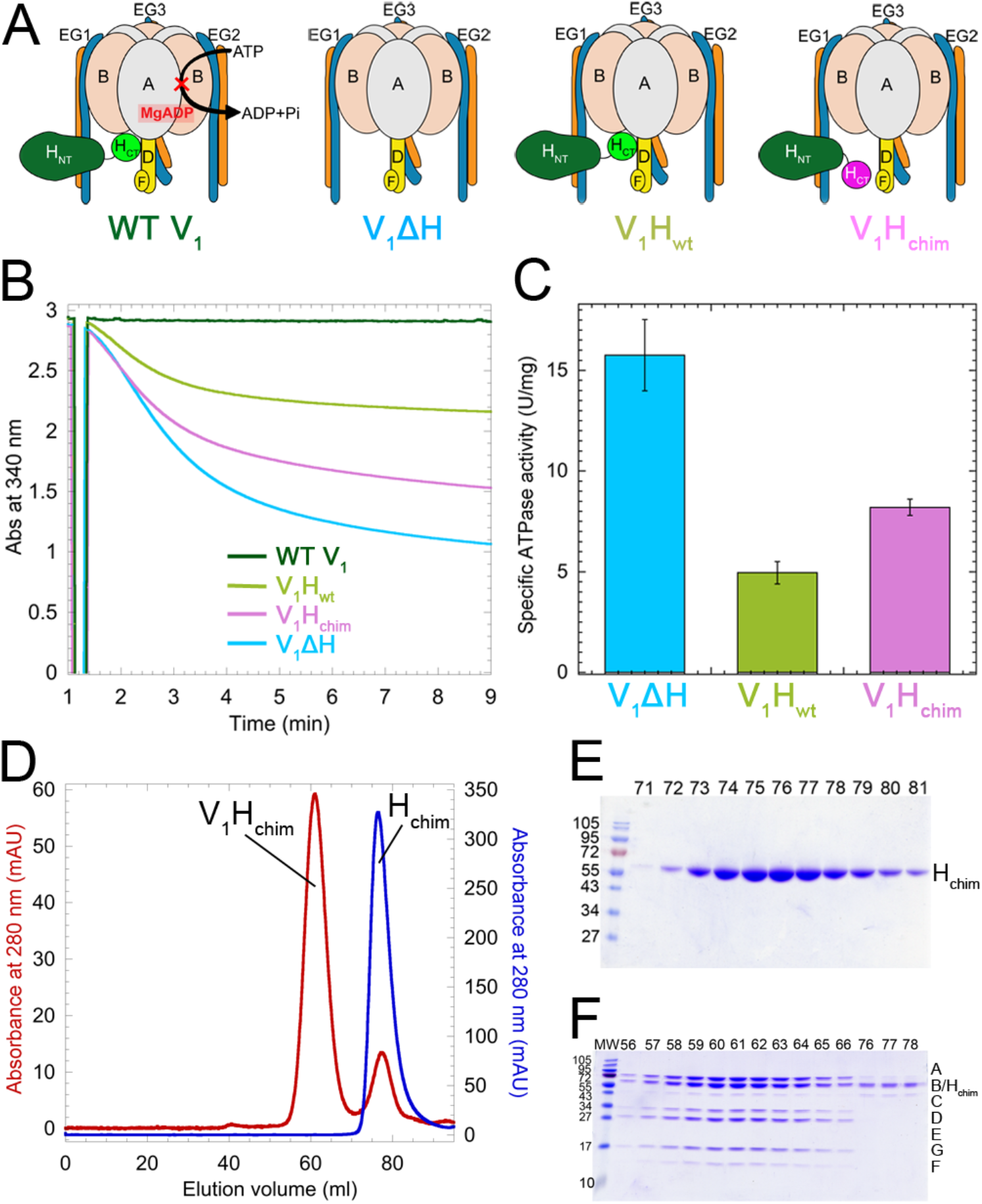
Purification and characterization of V_1_ mutants. A Schematic representation of the V_1_ mutants. B Time dependent MgATPase activities of the V_1_ mutants measured in an ATP regenerating assay. C Specific activities of the V_1_ mutants ± s.e.m from at least two independent purifications per mutant. D Size exclusion chromatography of H_chim_ (blue trace) and V_1_ΔH reconstituted with H_chim_ (V_1_H_chim_, red trace). E,F SDS-PAGE of column fractions of H_chim_ (E) and V_1_H_chim_ (F).

### V_1_H_chim_ and C subunit associate with V_o_ND to form coupled V-ATPase *in vitro*

The ability of above described V_1_ mutants to bind to V_o_ was tested using BLI. V_o_ND reconstituted with biotinylated MSP was immobilized on streptavidin coated BLI sensors (**Fig. 3*A***, step 1) and the V_o_ND coated sensors were then dipped in wells containing V_1_ mutants and subunit C (**Fig. 3*A***, step 2). We found that out of the four V_1_ mutants, only V_1_H_chim_ showed significant association with V_o_ND (**Fig. 3*A***, red trace). The BLI signal obtained was attributed to interactions at the V_1_-V_o_ interface as the observed on rate (kobs) was significantly (~ 60 fold) reduced in presence of a monoclonal antibody (10D7) against aNT, which is known to only bind free V_o_ (12) (**Supplementary Fig. S2**). The sensors were then dipped in buffer to measure dissociation rates (**Fig. 3*A***, step 3). However, no significant dissociation was observed, indicating stable assembly of V_1_H_chim_ with V_o_. Without C, none of the V_1_ mutants showed significant binding (**Supplementary Fig. S3**), consistent with studies in yeast that showed that deletion of C prevents assembly of V_1_V_o_ (36). As a control, the sensors were then dipped in prescission protease to cleave and release any remaining complex (**Fig. 3*A***, step 4).

**Fig. 3.**
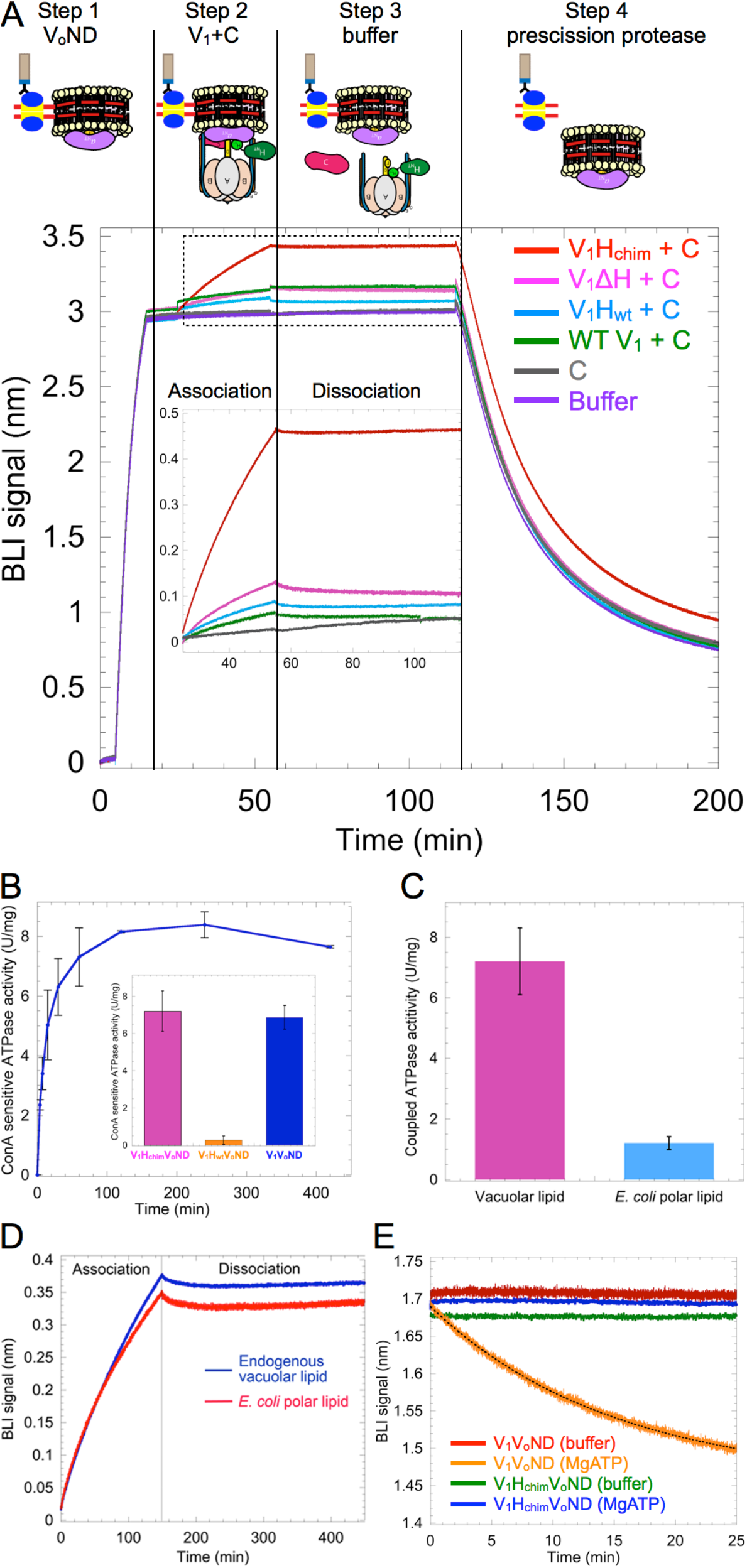
V_1_H_chim_ and C associate with V_o_ND to form coupled V_1_V_o_-ATPase. A V_o_ND was immobilized on streptavidin coated BLI sensors via biotinylated MSP (step 1). Sensors were then dipped into 0.4 μM of V_1_ mutants in presence of 1 μM C (association; step 2) followed by buffer (dissociation; step 3). Association with V_o_ND was most efficient with V_1_H_chim_ (red trace). Sensors were then dipped in prescission protease to verify that the BLI signal was not due to nonspecific binding (step 4). Inset shows an enlarged view of the association and dissociation steps B Equimolar amounts of V_1_H_chim_ and V_o_ND, and a twofold molar excess of C subunit were incubated at 22 °C, and the ConA sensitive MgATPase activity was measured as a function of time. Each point represents the mean ± s.e.m. of two separate reconstitutions from two individual purifications. Inset, specific MgATPase activities of reconstituted V_1_H_chim_V_o_ND and V_1_H_wt_V_o_ND (± s.e.m. from two independent purifications) compared to purified V_1_V_o_ND (34). C ConA sensitive MgATPase activity of V_1_H_chim_ reconstituted with V_o_ND containing vacuolar (pink) or *E. coli* polar lipid extract (blue). Activity data are from two separate reconstitutions using protein from two independent purifications. *(D)* Association and dissociation of V_1_H_chim_ with V_o_ND reconstituted with endogenous vacuolar lipids (blue trace) or *E. coli* polar lipids (red trace). One representative from two independent experiments is shown. E Following association of the V_1_H_chim_V_o_ND complex, sensors were dipped in wells containing buffer (green), or buffer + 1 mM MgATP (blue) for dissociation rate measurement. The dissociation phase of wild type V_1_V_o_ND in buffer (red) and buffer + 1 mM MgATP (orange) is included for comparison (data from (34)).

While we observed slow, but stable, association of V_1_H_chim_ and C with V_o_ND, from the BLI experiment it was not clear whether active and coupled V-ATPase is formed under these conditions. To address this question, we monitored MgATPase activity that is sensitive to the V-ATPase specific inhibitor, Concanamycin A (ConA) of a 1:1:2 mixture of V_1_H_chim_, V_o_ND and C as a function of time (**Fig. 3*B***). The experiment demonstrated that binding of V_1_H_chim_ to V_o_ND in presence of C resulted in the formation of a coupled V_1_H_chim_V_o_ND complex, and that the reconstitution under these conditions was complete in ~2 hours, with a final specific activity of 7.2 ± 1.09 U/mg, similar to what is reported for purified wild type V_1_V_o_ND (6.9 ± 0.6 U/mg) (34) (**Fig. 3*B***, inset, pink and blue bars, respectively). A similar experiment conducted with V_o_ND, C and V_1_H_wt_ produced only 0.26 ±0.2 U/mg of coupled activity (**Fig. 3*B***, inset, orange bar). Therefore, V_1_H_wt_ was unable to reconstitute with V_o_ and C under these experimental conditions, consistent with earlier *in vitro* studies (33) and the real-time BLI experiments presented here (**Fig. 3*A***).

### Vacuolar lipids are essential for coupled ATPase activity of V_1_H_chim_V_o_ND

Lipid moieties, such as long chain sphingolipids (37, 38), phosphoinositide PI(3,5)P_2_ (39), and ergosterol (40) have been implicated in V-ATPase assembly and function, but a direct involvement of these lipids in V-ATPase activity has not been shown. Since we were able to reconstitute the V_o_ sector in endogenous, vacuolar lipid containing nanodiscs, we wished to determine if vacuolar lipids played a role in the reconstitution of V_1_Hchim and C with V_o_ND. Similar to the experiment shown in **Fig. 3*B***, V_1_H_chim_ and C subunit were incubated with V_o_ND containing either endogenous vacuolar lipid, or *E. coli* polar lipid extract. We found significantly reduced coupled MgATPase activity of reconstituted V_1_H_chim_V_o_ND in *E. coli* polar lipid extract compared to endogenous vacuolar lipids (**Fig. 3*C***). To test if the lower activity was due to reduced binding of V_1_H_chim_ to *E. coli* lipid containing V_o_ND, we conducted a BLI experiment like the one described in **Fig. 3*A***. Endogenous vacuolar lipid or *E. coli* polar lipid containing V_o_ND was immobilized on streptavidin sensors using biotinylated MSP. The sensors were then dipped in V_1_H_chim_ and C subunit followed by buffer to assess formation and stability of V_1_H_chim_V_o_ND (**Fig. 3*D***). No significant difference was observed in the association or dissociation behavior between the two V_o_ND preparations. Therefore, while V_1_H_chim_ and C was able to bind V_o_ND containing *E. coli* polar lipids, endogenous vacuolar lipids were essential for the formation of coupled ATPase activity.

### V_1_H_chim_V_o_ND is more stable in presence of MgATP compared to V_1_V_o_ND

Using BLI, we have previously demonstrated that V_1_’s dissociation from V_o_ is negligible under non-ATP hydrolyzing conditions, but that in presence of MgATP, the complex undergoes spontaneous dissociation at a rate of 1 x 10^−3^ ± 3.3 x 10^−6^ s^−1^ (34). We conducted a similar experiment using V_1_H_chim_V_o_ND, wherein after association of V_1_H_chim_ and C with V_o_ND on BLI sensors, we dipped the sensors in wells containing buffer or 1 mM MgATP. Unlike V_1_V_o_ND, V_1_H_chim_VoND showed negligible dissociation in the presence of MgATP, suggesting that the assembled V_1_H_chim_V_o_ND complex is inherently more stable than V_1_V_o_ND (**Fig. 3*E***). The experiment thus highlights the importance of H_CT_’s conformational switch in driving V-ATPase disassembly.

### Structural and functional characterization of V_1_H_chim_V_o_ND

To determine the efficiency of V_1_H_chim_V_o_ND complex formation, equimolar amounts of V_1_H_chim_ and V_o_ND with a 2-fold excess of C were incubated for up to 16 hours at 22 °C, and the reconstitution mixture was resolved by glycerol density gradient centrifugation. SDS-PAGE of the gradient fractions showed that the majority of V_1_ and V_o_ subunits co-migrated to fractions 7-10, similar to what was observed for purified wild type V_1_V_o_ND (34), with the excess C subunit remaining in lighter fractions (**Fig. 4*A***). Negative stain EM of the peak fractions showed single particles of V_1_H_chim_V_o_ND, with the typical dumbbell shaped appearance of V_1_V_o_ observed in multiple studies (26, 28, 34, 41, 42) (**Fig. 4*B***). A more detailed analysis indicated a good match between averages obtained by reference free alignment and classification of a small dataset of V_1_H_chim_V_o_ND and corresponding projections of a cryoEM model of yeast V_1_V_o_ (28) (**Fig. 4*C***).

**Fig. 4.**
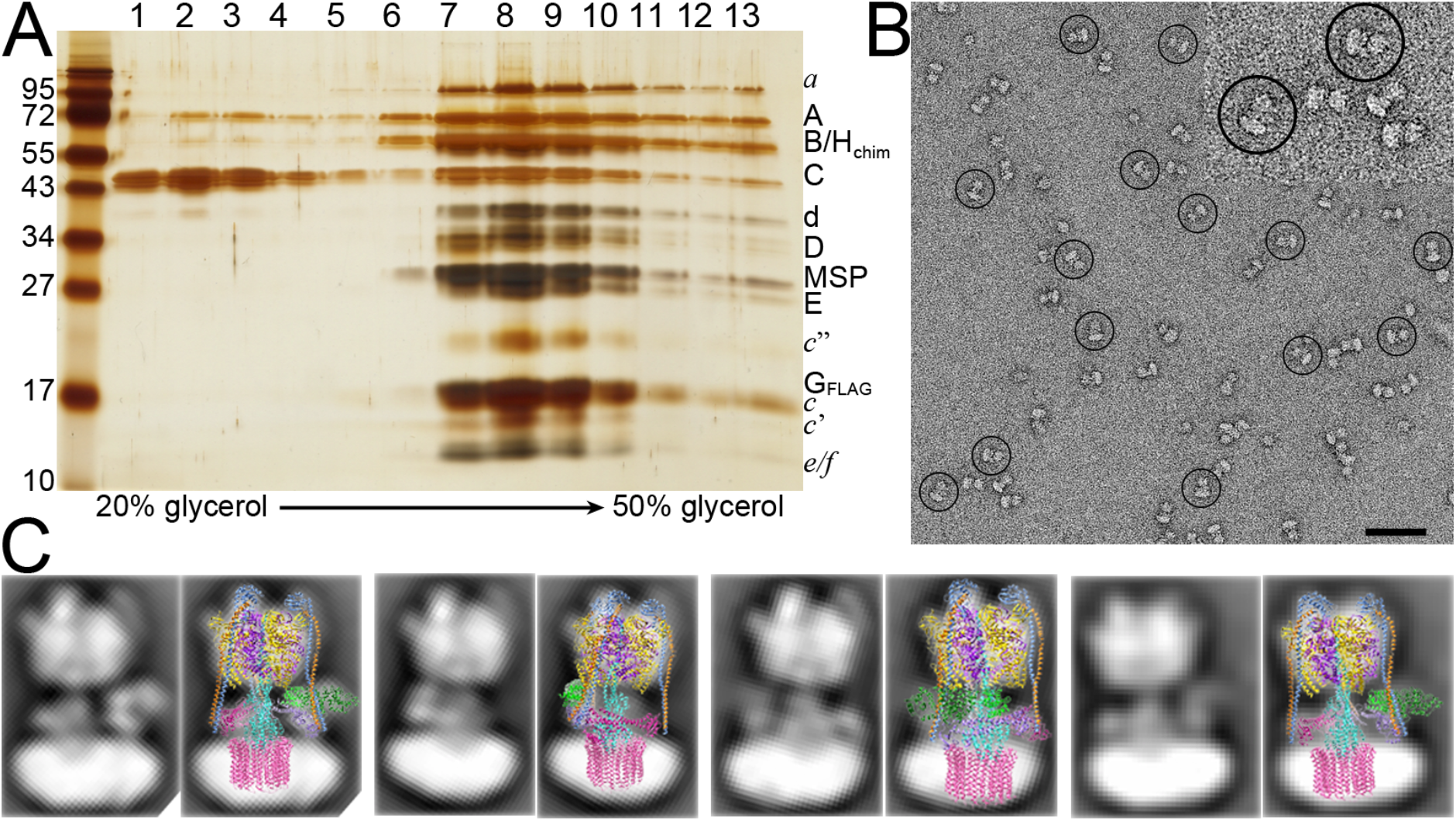
Structural and functional characterization of the V_1_H_chim_V_o_ND complex. A Reconstituted V_1_H_chim_V_o_ND was subjected to glycerol gradient centrifugation and the gradient fractions were analyzed by silver stained SDS-PAGE. B Negative stain EM of V_1_H_chim_V_o_ND showing homogeneous and monodisperse dumbbell shaped molecules. C A dataset of ~5800 particle projections was subjected to reference-free alignment and classification, and selected class averages were overlaid with projections of the cryoEM model of yeast V_1_V_o_ (28) (pdb id: 3J9U). Bar in (B) = 50 nm.

## Discussion

### The release of H_CT_ from its inhibitory position is necessary and sufficient for *in vitro* reconstitution of V_1_ and C with V_o_

In our experiments, only V_1_H_chim_ shows significant binding to V_o_ in the presence of C. As mentioned earlier, in V_1_H_chim_, H_CT_ lacks the inhibitory loop (32) (**Supplementary Fig. S1*A-D***, red spheres), which probably reduces H_CT_’s affinity for V_1_, in turn diminishing H subunit mediated inhibition of V_1_’s MgATPase activity (**Fig. 2*B,C***). We therefore conclude that for V_1_ to reconstitute with V_o_, H_CT_ must be released from its inhibitory position on V_1_, making it available for binding to *a*_NT_ as part of V_o_. Upon disassembly of V_1_ from V_o_, *a*_NT_ moves from a peripheral position in V_1_Vo to a more medial position, where it binds subunit *d* in autoinhibited V_o_ (**Supplementary Fig. S1*E-G***) Four separate EM reconstructions of purified yeast V_o_ show *a*_NT_ in the medial position bound to *d* (16, 29–31) and so we conclude that in our preparation of V_o_ND, *a*_NT_ is in a similar autoinhibitory conformation (**Supplementary Fig. S1*E-G***) In our reconstitution experiments, V_1_H_chim_, C subunit and autoinhibited V_o_ND are sufficient to form a structurally and functionally coupled V-ATPase, suggesting that no conformational change in *a*NT is required, and the release of H_CT_ from its autoinhibitory position on V_1_ is necessary and sufficient for reassembly of V_1_ and V_o_. Our finding is consistent with the fact that reassembly of V_1_ with V_o_ on vacuoles is not inhibited by Con A, an inhibitor of c-ring rotation in the V_o_ sector (14).

### Release of inhibitory MgADP may cause detachment of H_CT_ from its inhibitory position on V_1_

H_NT_ and H_CT_ occupy specific binding sites on free V_1_, with H_NT_ bound to EG1 and H_CT_ bound to the bottom of the A_3_B_3_ hexamer, with its inhibitory loop wedged between the B subunit of an open catalytic site and the central stalk (**Supplementary Fig. S1*A-D***) (32). The specific interaction of H_CT_ with an open catalytic site maintains inhibitory MgADP in the adjacent closed catalytic site, locking autoinhibited V_1_ in rotational state 2. We have recently observed that transient MgATP hydrolysis on V_1_H_wt_, which, unlike wild type V_1_, is not in the MgADP inhibited state (**Fig. 2*B***), lowers H’s affinity for V_1_, and we reasoned that this destabilization of the V_1_-H interaction is caused by MgATP hydrolysis driven conformational changes at the catalytic sites and the central (DF) and peripheral stalks (EG1-3) (24). We propose that an allosteric structural change at the open catalytic site driven by release of inhibitory MgADP from the closed catalytic site in autoinhibited, wild type V_1_ (by a yet unknown mechanism) leads to the detachment of H_CT_ from its inhibitory position at the open site, making it in turn available to bind *a*_NT_ on V_o_.

### Incorporation of C subunit and the possible role of the RAVE complex

From studies in yeast it was shown that while C is required for binding of V_1_ to V_o_ on yeast vacuolar membranes (36), deletion of H allows assembly of a labile, but inactive, complex (27, 43). The requirement of C for association of V_1_ with V_o_ is supported by our *in vitro* BLI experiments, wherein none of the V_1_ mutants reconstituted with V_o_ in absence of C (**Supplementary Fig. S3**). However, unlike *in vivo*, the presence of C along with V_1_ΔH, wild type V_1_, or V_1_H_wt_ is not sufficient for reconstituting V_1_V_o_ *in vitro*. This discrepancy is not due to a reduced affinity of C for V_1_, as we obtained a K_d_ of ~0.7 nM for the interaction between C and V_1_ΔC (**Supplementary Fig. S4**). A high-affinity interaction between C and V_1_ΔC is consistent with sub-stoichiometric amounts of C remaining associated with purified V_1_ΔH and V_1_ (32), as well as the reported K_d_ of ~42 nM for the EG-C interaction (25). However, tight binding of C to V_1_ΔC is inconsistent with the release of C into the cytosol upon disassembly of V_1_ from V_o_ in yeast. It has been reported that in the presence of the microtubule depolymerizing drug benomyl, C does not dissociate from V_1_V_o_ upon glucose removal (44). In addition, a direct interaction between C and tubulin has been observed (44, 45), suggesting the possibility that, upon disassembly, C is sequestered by microtubules, preventing its re-association with V_1_ and/or V_o_.

While incorporation of C upon glucose addition occurs in absence of microtubules (44), efficient (re)assembly of holo V-ATPase requires a heterotrimeric chaperone complex referred to as regulator of H^+^-ATPase of vacuolar and endosomal membranes (RAVE). It has been proposed that upon receiving the signal for reassembly, RAVE recruits C and V_1_ to V_o_ on vacuolar membranes by directly interacting with C, EG (as part of V1) and *a*_NT_ (as part of V_o_) (46). Under the *in vitro* conditions employed here, it takes ~2 hours for a 1:1:2 mixture of V_1_H_chim_, V_o_ND and C to complete reconstitution of V_1_H_chim_V_o_ND, a relatively slow process compared to the kinetics of reassembly observed *in vivo* (~5 minutes (47)). It is possible that the RAVE complex, by increasing the proximity of V_1_, C and V_o_, facilitates the otherwise low affinity interactions of the V_1_-V_o_ interface, thereby accelerating reassembly.

### Proposed model for the reassembly of V_1_ and C with V_o_ on the vacuolar membranes of yeast

Based on the here presented data, we conclude that the detachment of HCT from V_1_ and the presence of C subunit are required for the reassembly of V_1_ with V_o_ (**Fig. 5*A-C***). In our *in vitro* reconstitutions, the association between V_1_H_chim_ and V_o_ND is driven by the H_CT_-*a*_NT_ interaction (**Fig. 5*B***), but *in vivo*, the chain of events that leads to reassembly of autoinhibited V_1_ and V_o_ are probably different, as H_CT_ is in its inhibitory conformation on V_1_ (**Fig. 5*A***). We propose that in yeast, upon receiving cellular signals, autoinhibited V_1_ and C are first recruited to V_o_ (**Fig. 5*D***), a process that is likely the rate limiting step for reassembly. Our reason for this hypothesis is that even with the requirements for reassembly being met in our *in vitro* reconstitution of V_1_H_chim_ and C with V_o_, the rate of reassembly was slow. *In vivo*, recruitment of V_1_ and C to V_o_ is facilitated and probably accelerated by the RAVE complex (46), but efficient (re) assembly *in vivo* requires additional factors such as the glycolytic enzymes aldolase (48) and phosphofructokinase (47), whose function in the process is currently not known. Once V_1_ and C are recruited to V_o_ at the vacuolar membrane, inhibitory MgADP is released upon unclasping of the closed catalytic site by a yet unknown mechanism (**Fig. 5*E***). The release of inhibitory MgADP allows MgATP hydrolysis to resume, with concomitant conformational changes at the catalytic sites and rotation of the central stalk (DF), structural changes that result in detachment of H_CT_ from V_1_ (24) (**Fig. 5*F***). The proximity of *a*_NT_ to V_1_-detached H_CT_ facilitates the H_CT_-*a*_NT_ interaction, a requirement for coupling of V_1_ to V_o_ in the holo-enzyme (27). The H_CT_-*a*_NT_ interaction stabilizes the peripheral conformation of *a*_NT_ such that the C_foot_-EG2-*a*_NT_ ternary complex can be formed, thus completing functional (re)assembly (**Fig. 5*C***).

**Fig. 5.**
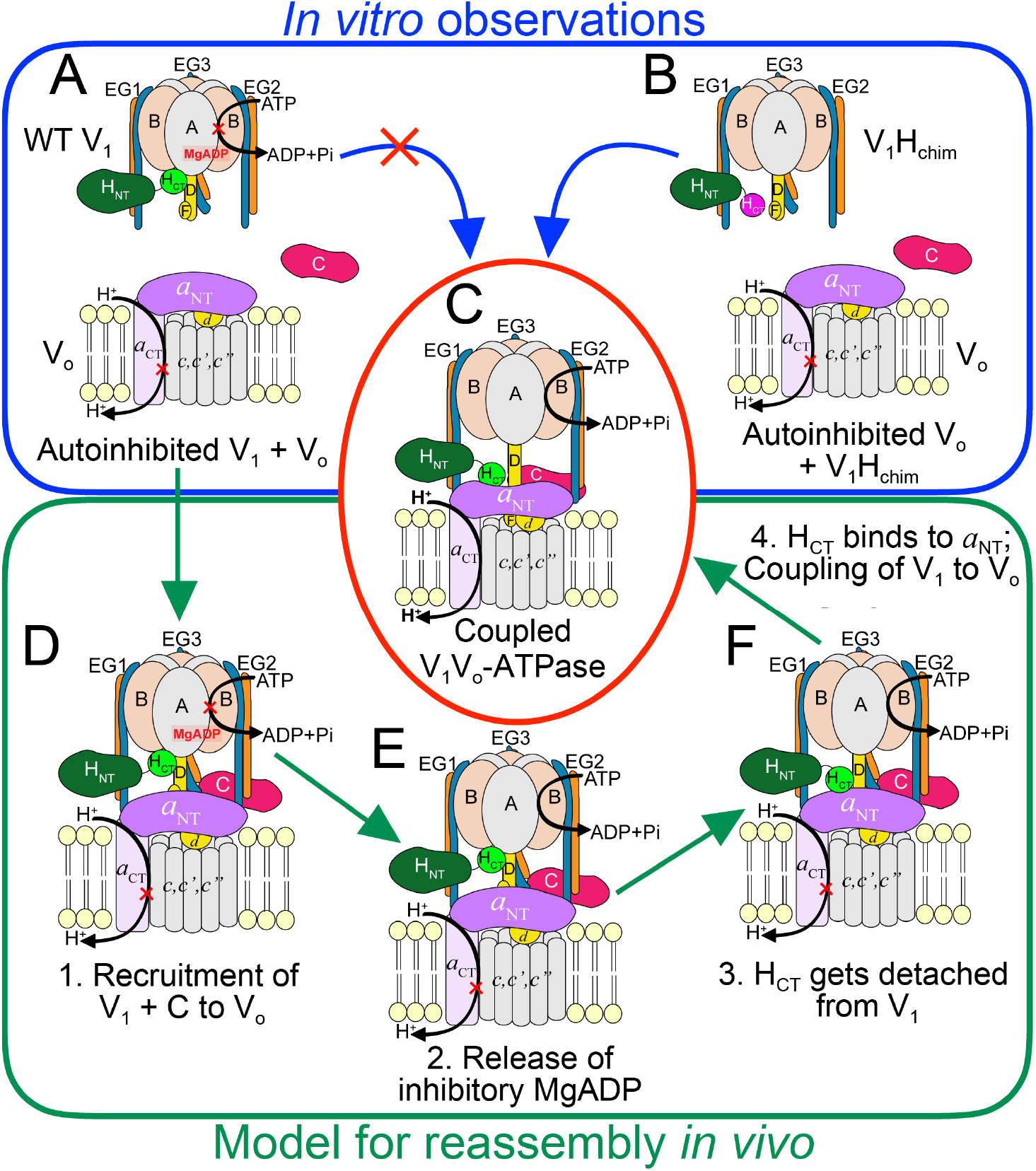
Model for reassembly of autoinhibited V_1_ and V_o_. A-C Our *in vitro* experiments have shown that while wild type H containing V_1_ does not readily bind V_o_ND (A), V_1_H_chim_ spontaneously associates with V_o_ND (B) to form a structurally and functionally coupled V-ATPase, albeit at a slow rate (C). *In vivo*, however, V_1_ exists in the autoinhibited conformation (A), and the rate of assembly with V_o_ is significantly faster (within 5 minutes). D-F For *in vivo* (re)assembly, we propose that the following steps occur: 1., recruitment of V_1_ and subunit C to the vacuolar membrane (D); 2., release of inhibitory MgADP (E); 3., detachment of H_CT_ from its inhibitory position on V_1_ and 4., H_CT_ binding to *a*_NT_ (F). For further details, see text.

## Materials and Methods

### Strains

The yeast strain SF838-5Aα deleted for the vma2 gene (B subunit) vma2Δ::Nat was a kind gift from Dr. Patricia Kane, SUNY Upstate Medical University. A plasmid containing the FLAG tag with a KanMX6 marker, pFA6a-6xGly-FLAG-kanMX6 was a gift from Dr. Mark Hochstrasser (49) (Addgene plasmid #20751). The primers, vph1CTFlagFWD:gctgttgctagtgcaagctcttccgcttcaagcGGGGGAGGCGGGGGTGGAA and vph1CTFlagREV: cctggatgtggatttcgattctaacgttaccccaaggcaaatgatggtcactggtgg GAATTCGAGCTCGTTTAAAC were used to amplify the FLAG tag and KanMX marker from pFA6a-6xGly-FLAG-kanMX6. The ~ 1.8 kb product was gel purified and used for homologous recombination to insert the FLAG-KanMX cassette in the C terminus of vph1 in the yeast strain SF838-5Aα vma2Δ::Nat using the same primers as above. Colonies were selected for growth on YPD G418 plates and the insertion of the FLAG tag at the C terminus of vph1 was confirmed by sequencing. The construction of chimeric H subunit (H_chim_) encoding the N terminal domain from *S. cerevisiae* (1-352) and the C-terminal domain (349-483) of the human H subunit into the yeast pRS316 vector has been discussed in (32). From the pRS316 vector, using the primers MalChimF: TTAGCCGGTACCGGGAGCAACGAAGATATTAATGGAC and MalChimR: TTACCAAAG CTTTTAGCTTCGGGCGGCAG, H_chim_ was amplified. The primers additionally introduced the restriction sites 5’ KpnI and 3’ HindIII which were used insert the amplicon into a pMal vector. The resultant vector encoded MBP tagged H_chim_ separated by a Prescission protease cleavage site, as confirmed by sequencing.

### Purification of V_o_ and its reconstitution into endogenous vacuolar lipid

V_o_ was purified from yeast vacuoles and reconstituted into endogenous vacuolar lipid containing nanodiscs as described for V_1_V_o_ND in (34). The steps are briefly described as follows.

### Purification of biotinylated membrane scaffold protein (MSP)

Biotinylated MSP was purified as described in (34). Briefly, BL21 (DE3) cells were co-transformed with the plasmids pHBPMSP1E3D1 and pBirAcm (encoding the BirA gene). Cells were grown in rich broth supplemented with 0.1 mM D-biotin, 34 μg/ml chloramphenicol and 30 μg/ml kanamycin to an OD_595_ of ~ 0.5 at 37 °C followed by induction using 0.5 mM IPTG for 3 to 4 hours. Harvested cells were purified as described in (30). Briefly, cells were lysed by sonicating for 3 x 30 seconds. The lysate was cleared by centrifugation at 13,000 x *g* and passed over a nickel-nitrilotriacetic acid affinity column. The column was washed with 10 column volumes of the each of the three buffers: 40 mM Tris-HCl, 300 mM NaCl and 1 % Triton X-100 pH 8; 40 mM Tris-HCl, 300 mM NaCl, 50 mM sodium cholate and 5 mM immidazole, pH 8; 40 mM Tris-HCl, 300 mM NaCl and 10 mM immidazole, pH 8. MSP was eluted with a 10 column volume gradient of the elution buffer (40 mM Tris-HCl, 300 mM NaCl and 100 mM immidazole, pH 8). Purified biotinylated MSP was dialyzed into 25 mM Tris, 150 mM NaCl, 0.5 mM EDTA, pH 7.2, concentrated to ~ 5 mg/ml, snap frozen in liquid nitrogen and stored at – 80 °C until use.

### Isolation of yeast vacuoles

Yeast vacuoles were isolated by flotation on a Ficoll density gradient as described in (50). Briefly, SF838-5Aα vma2Δ::Nat with a Flag tag on the C terminus of vph1 (*a* subunit) was grown to an OD_595_ of ~ 1.0 in YPD pH 5. 12 liters of cells were harvested by centrifugation at 5000 x *g* for 30 minutes. The pellet was washed and resuspended in 100 ml of 1.2 M sorbitol with ~ 15 mg of zymolyase to form spheroplasts. The spheroplasts were recovered in 100 ml each of 2.4 M sorbitol and 2x YPD and then resuspended in buffer containing 12 % Ficoll 400. The suspension was homogenized in a dounce homogenizer and centrifuged at 71,000 x *g* for 40 minutes. Vacuole wafers from the top of the gradient were extracted, homogenized in buffer containing 8 % Ficoll and centrifuged at 71,000 x *g* for 40 minutes. The final vacuole wafers were resuspended in 1.5 mM Mes-Tris, pH 7.0, 5 % glycerol and 1 mM β-mercaptoethanol. Vacuolar protein concentration was measured using a modified BCA assay (16) and the vacuoles were frozen in liquid nitrogen until further use.

### Extraction of V_o_ and reconstitution into lipid nanodiscs

Three batches of purified vacuoles (12 liters each) were typically used for one extraction as described in (34). Briefly, thawed vacuoles were combined, supplemented with protease inhibitors and solubilized using 1.2 mg of n-dodecyl β-D-maltopyranoside (DDM) per 1 mg of vacuolar protein. To the detergent solubilized sample, purified biotinylated MSP was added in a molar ratio of 1:50 (vacuolar protein: MSP). The mixture containing vacuolar protein, vacuolar lipids and MSP was incubated at 4 °C for 1 hour followed by detergent removal using bio-beads. Reconstituted vacuolar membrane proteins in biotinylated and endogenous vacuolar lipid containing nanodiscs were subjected to anti-FLAG affinity chromatography to purify V_o_ containing nanodiscs. The eluate from the FLAG column was then subjected to size exclusion chromatography using a Superdex 200 1 x 30 cm column. Peak fractions from gel filtration were combined and concentrated using a Vivaspin 100,000 MWCO concentrator.

### Purification of the chimeric H subunit (Hchim)

*E. coli* Rosetta2 (Novagen) cells expressing N-terminal MBP tagged H_chim_ were grown to an OD_600_ ~0.5 (in LB, 0.2% glucose, 50 μg/ml carbenicillin and 34 μg/ ml chloramphenicol) and expression was induced with 0.5 mM IPTG at 30 °C for 4 hours. Protein was purified using amylose affinity chromatography and the MBP tag cleaved with Prescission protease as previously described (25). The pH of the cleavage product was adjusted to 7 by overnight dialysis in 25 mM sodium phosphate pH 7, 0.5 mM EDTA and 5 mM BME. At pH 7, H_chim_ has a predicted charge of + 3.5 whereas the predicted charge of MBP is −9 (Protein Calculator v3.4), allowing separation of the two proteins using cation exchange (carboxymethyl, CM) chromatography. The cleaved MBP came off the CM column in the flow through and wash steps, while pure H_chim_ was eluted in dialysis buffer supplemented with 100 mM NaCl. The preparation was subjected to a final step using a Superdex 200 1.6 x 50 cm size exclusion chromatography column.

### Purification of V_1_ mutants and subunit C

WT V_1_ and V_1_ΔH were purified from vma10Δ::KanMX (34) and vma13Δ::KanMX, vma10Δ::Nat (35) respectively as described in (32). In both cases, yeast cells were transformed with a pRS315 vector containing N-terminally FLAG tagged vma10 (G subunit) (51). Cells were grown to an OD_595_ of ~ 4 in synthetic dropout medium without Leucine (SD-Leu) and harvested by centrifugation at 4000 x *g* for 15 minutes. Cells were lysed by ~ 15 passes through a microfluidizer (Microfluidics M-110L). Unbroken cells were pelleted by centrifugation at 4000 x *g* for 30 minutes and the resultant supernatant was cleared by centrifugation at 13,000 x *g* for 40 minutes. Cleared lysate was subjected to affinity chromatography using anti-FLAG resin. The eluate from the FLAG column was concentrated and subjected to size exclusion chromatography using a Superdex 200 1.6 x 50 cm column.

Purified V_1_ΔH was incubated for one hour at 4 °C with a ~ 5 fold molar excess of either Hwt (purified as in (24)) or H_chim_ to form the V_1_H_WT_ and V_1_H_chim_ complex respectively. Excess H_WT_ or H_chim_ was removed by gel filtration using a Superdex 200 1.6 x 50 cm column. Subunit C was purified as described in (52).

### Purification of V_o_ from yeast membranes and its reconstitution into E. coli lipid containing nanodiscs

The purification of V_o_ from yeast membranes was adapted from (16). SF838-5Aα vma2Δ::Nat with a FLAG tag on the C-terminus of vph1 was grown in YPD pH 5 media to an OD595 of ~ 8 and harvested by centrifugation at 4000 x *g* for fifteen minutes. Cell pellets were resuspended in lysis buffer (TBSE buffer supplemented with 8 % sucrose, 2 % sorbitol and 2 % glucose) and stored at - 80 °C until use. To the thawed cell pellets, protease inhibitors were added to final concentrations of 2 μg/ml for leupeptin and pepstatin, 1 μg/ml chymostatin and 1 mM PMSF. Cells were lysed by bead-beating using an OMNI MACRO ES programmable digital homogenizer at 20,000 rpm with 0.5 mm Zirconia beads (BioSpec). The chamber was kept in an ice/salt slush to maintain low temperature during lysis. Cell debris and mitochondria were removed by centrifugation at 1200 x *g* for 10 minutes followed by a centrifugation at 21,000 x *g* for 20 minutes. Crude membranes were isolated by centrifugation at 370,000 x *g* for 1 hour using a Ti-70 rotor in a Beckman ultracentrifuge. The pellet was gently resuspended in membrane wash buffer (25 mM Tris, pH 7.2, 0.5 M Sorbitol), homogenized in one stroke and centrifuged again at 370,000 x *g* for 1 hour. The final membrane pellet obtained was resuspended in membrane wash buffer and stored at - 80 °C until use. Thawed membranes were supplemented with protease inhibitors (2 μg/ml for leupeptin and pepstatin, 1 μg/ml chymostatin and 1 mM PMSF) and solubilized using 1 mg of n-dodecyl β-D-maltopyranoside (DDM) per mg of membrane protein. The suspension was rotated at 4 °C for 1 hour and the insoluble fraction was pelleted by centrifugation at 180,000 x *g* for 1 hour. The supernatant was subjected to affinity chromatography using a 3 ml FLAG column. The peak fractions from the FLAG column was reconstituted into lipid nanodiscs using a molar ratio of 0.02 V_o_: 1 MSP: 25 lipid (30). 1.25 % DDM (w.r.t the final volume of the reconstitution mixture) was added to *E. coli* polar lipid extract followed by V_o_ and biotinylated MSP. The reconstitution mixture was rotated at 4 °C for 1 hour and the detergent was removed using ~ 0.5 g of activated bio-beads for 2 hours. V_o_ reconstituted in lipid nanodiscs were separated from empty nanodiscs using size exclusion chromatography (Superdex 200 1.6 x 50 cm column).

### Biolayer interferometry (BLI)

Interaction of V_o_ with the purified V_1_ mutants was screened using BLI, a light interference based technique, similar to surface plasmon resonance. An Octet-RED system with streptavidin coated biosensors (FortéBio, SA biosensors, catalog number 18-5019) were used for the experiments. All BLI experiments were conducted using 25 mM Tris-HCl pH 7.2, 150 mM NaCl, 0.5 mM EDTA, 1 mM BME, 0.5 mg/ml BSA. The temperature was maintained at 22 °C, with each biosensor stirred in 0.2 ml of sample at 1000 rpm and a standard measurement rate of 5 s^−1^. Streptavidin coated biosensors were pre-wetted in BLI buffer and then dipped in wells containing 3 μg/ml of biotinylated V_o_ND. A buffer control was included to show that none of the buffer components interacted with the sensors. Details of individual experiments have been described in the respective figure legends. The affinity of interaction between V_1_ΔC and MBP-C was measured using anti-mouse IgG Fc capturing biosensors (FortéBio, AMC biosensors catalog number 18-5088) as described (24).

### ATPase activity assay

MgATPase activity of purified V_1_ mutants and reconstitution mixtures (V_1_ mutants + V_o_ND + subunit C) was measured using a coupled enzyme assay as described in (32). Briefly, 10 μg of the V_1_ mutant was added to an assay mixture containing 1 mM MgCl2, 5 mM ATP, 30 units/ml each of lactate dehydrogenase and pyruvate kinase, 0.5 mM NADH, 2 mM phosphoenol pyruvate, 50 mM HEPES, pH 7.5, at 37 °C. The decrease of absorbance at 340 nm corresponding to the decline of NADH in the system was measured in the kinetics mode on a Varian Cary Bio100 spectrophotometer. In case of reconstitution mixtures, 20 μg of V_1_ mutant with equimolar amounts of V_o_ND and a 2X molar excess of C subunit was added to an assay containing 4 mM MgCl2.

## Author Contributions

Conceptualization, S.S., R.A.O. and S.W.; Investigation, S.S. and R.A.O.; Writing, S.S.; Editing, S.S., R.A.O. and S.W.; Funding Acquisition, S.W.

## Acknowledgements

We thank Dr. Patricia Kane for reagents and many helpful discussions and Dr. Thomas Duncan for his assistance with BLI data collection and analysis. This work was supported by NIH grant GM058600 to S.W. The content is solely the responsibility of the authors and does not necessarily represent the official views of the National Institutes of Health.

## Conflict of Interest

The authors declare that they have no conflicts of interest with the contents of this article.

## Abbreviations

V-ATPase, vacuolar H^+^-ATPase; V_1_, ATPase sector of the V-ATPase; V_o_, membrane sector of the V-ATPase; *a*_NT_, N-terminal cytoplasmic domain of the *a* subunit; *a*_CT_, C-terminal transmembrane domain of the *a* subunit; H_CT_, C-terminal domain of the H subunit; H_NT_, N-terminal domain of the H subunit; BME, β-mercaptoethanol; MBP, maltose binding protein; BLI, biolayer interferometry; TCEP, tris(2-carboxyethyl)phosphine; ConA, Concanamycin A.

## Supporting Information

**Supplementary Fig. S1.**
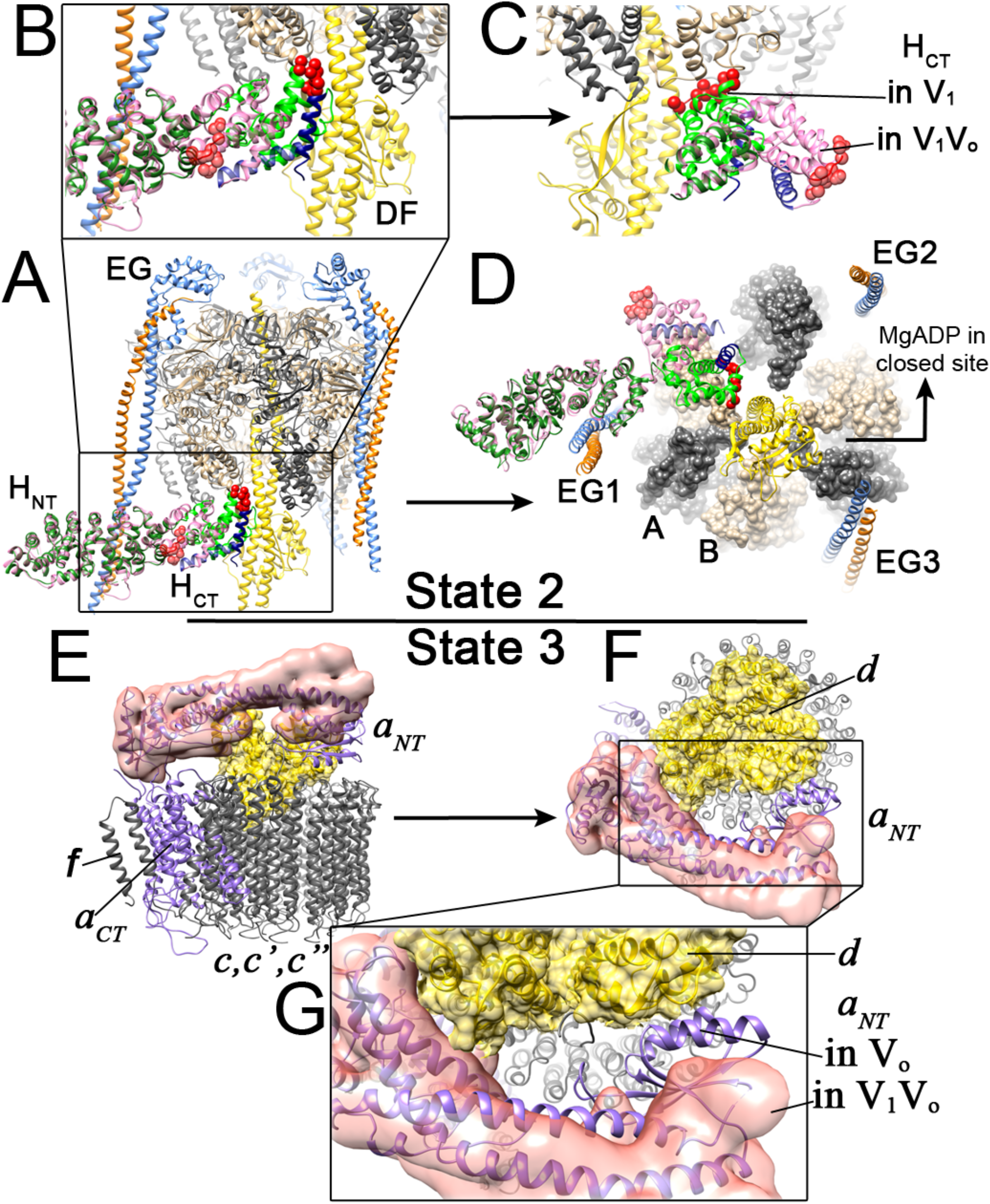
Structural changes in the V_1_-V_o_ interface subunits upon disassembly of V_1_ from V_o_. Out of the three rotational states of V_1_V_o_ (1), the crystal structure of autoinhibited V_1_ (2) aligns with state 2, while the cryo EM structure of autoinhibited V_o_ (3, 4) aligns with state 3, highlighting the conformational mismatch at the V_1_-V_o_ interface. *A*, Crystal structure of autoinhibited V_1_ sector showing the H subunit in its inhibitory conformation (green) (pdb id: 5D80) (2) overlaid with the conformation of H in V_1_V_o_ (pink) (pdb id: 3J9U) (1). *B*, Enlarged view of H subunit shows that H_CT_ undergoes a large conformational change from its conformation in V_1_V_o_ to autoinhibited V_1_. The terminal helix of H_CT_ is colored blue and the amino acids of the inhibitory loop are depicted as red spheres. *C*, A 90° rotated view of *B* across the y-axis. A comparison of the two conformations of H revealed that H_CT_ undergoes a 150° rotation from its conformation in V_1_V_o_ to that in V_1_. *D*, A 90° rotation of *A* across the x-axis shows that in V_1_, the inhibitory loop is wedged between the B subunit of an open catalytic site and the central stalk (DF) in turn promoting the entrapment MgADP in the adjacent closed catalytic site. *E*, CryoEM structure of autoinhibited V_o_ showing *a*_NT_ in its inhibitory conformation (purple ribbon) (pdb id: 6C6L (3)) overlaid with EM density for the conformation of *a*_NT_ in V_1_V_o_ (pink surface) (pdb id. 3J9V, (1). *F*, 90° rotation of *E* across the x-axis shows that *a*_NT_ undergoes a conformational change from a lateral position (pink) in V_1_V_o_ to a medial position in autoinhibited V_o_. *G*, Enlarged view of the highlighted region in *F* shows that in V_o_, *a*_NT_ interacts with *d*, potentially to close the proton pore.

**Supplementary Fig. S2.**
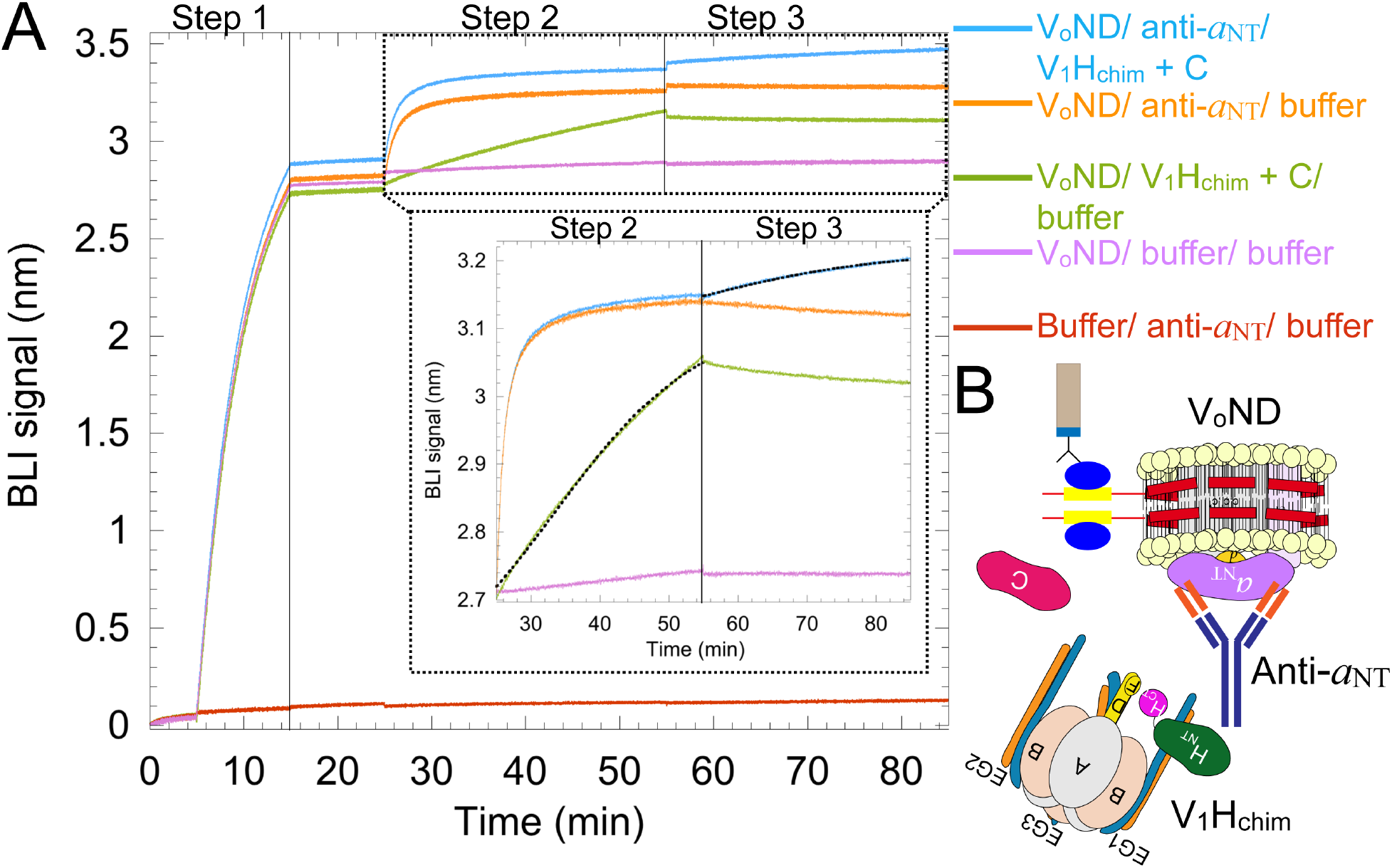
Assembly of V_1_H_chim_ with V_o_ is inhibited by an ant specific monoclonal antibody. *A*, V_o_ND was immobilized on streptavidin sensors and the sensors were then dipped in either α-*a*_NT_ mAb (10D7; blue and orange), V_1_H_chim_ + C (green) or buffer (pink) (step 2). V_1_H_chim_ + C bound to V_o_ND was then dipped in buffer while α-*a*_NT_ mAb bound to V_o_ND was dipped in V_1_H_chim_ + C (blue) or buffer (orange) (step 3). The red buffer trace was subtracted from all other traces and the resultant curves are shown in the inset. The curves obtained for V_1_H_chim_ + C’s association with V_o_ND in the presence (blue) or absence (green) of bound α-*a*_NT_ mAb were fit to a single site binding equation (dotted lines). Observed on rates (kobs) were 1.1 x 10^−3^ ± 2.7 x 10^−6^ without α-αNT mAb (green) and 7.3 x 10^−4^ ± 3.4 x 10^−6^ with α-*a*_NT_ mAb (blue). A representative experiment out of two repeats is shown here. *B*, Schematic showing inhibition of V_1_H_chim_ + C’s interaction with V_o_ND in presence of α-*a*_NT_ mAb bound to *a*_NT_ on V_o_.

**Supplementary Fig. S3.**
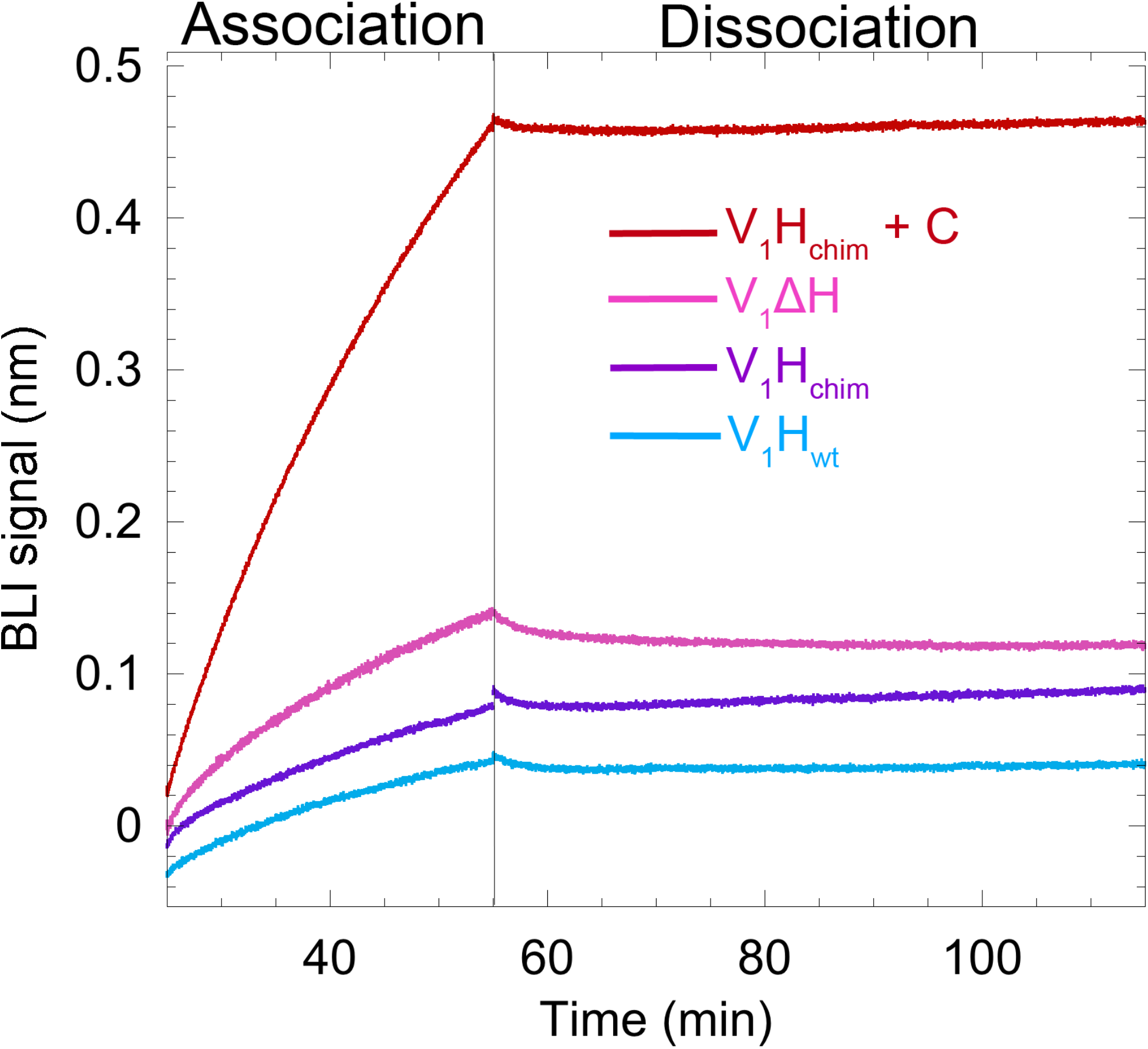
V_1_ mutants do not reconstitute with V_o_ in the absence of subunit C. V_o_ND immobilized on streptavidin sensors was dipped in 0.4 μM V_1_ΔH (pink), V_1_H_wt_ (blue), V_1_H_chim_ (purple) and V1H_chim_ with 1μM C subunit (red) followed by buffer. Compared to V_1_H_chim_ + C, none of the V_1_ mutants showed significant signal for binding to V_o_ND. A representative from two separate experiments is shown.

**Supplementary Fig. S4.**
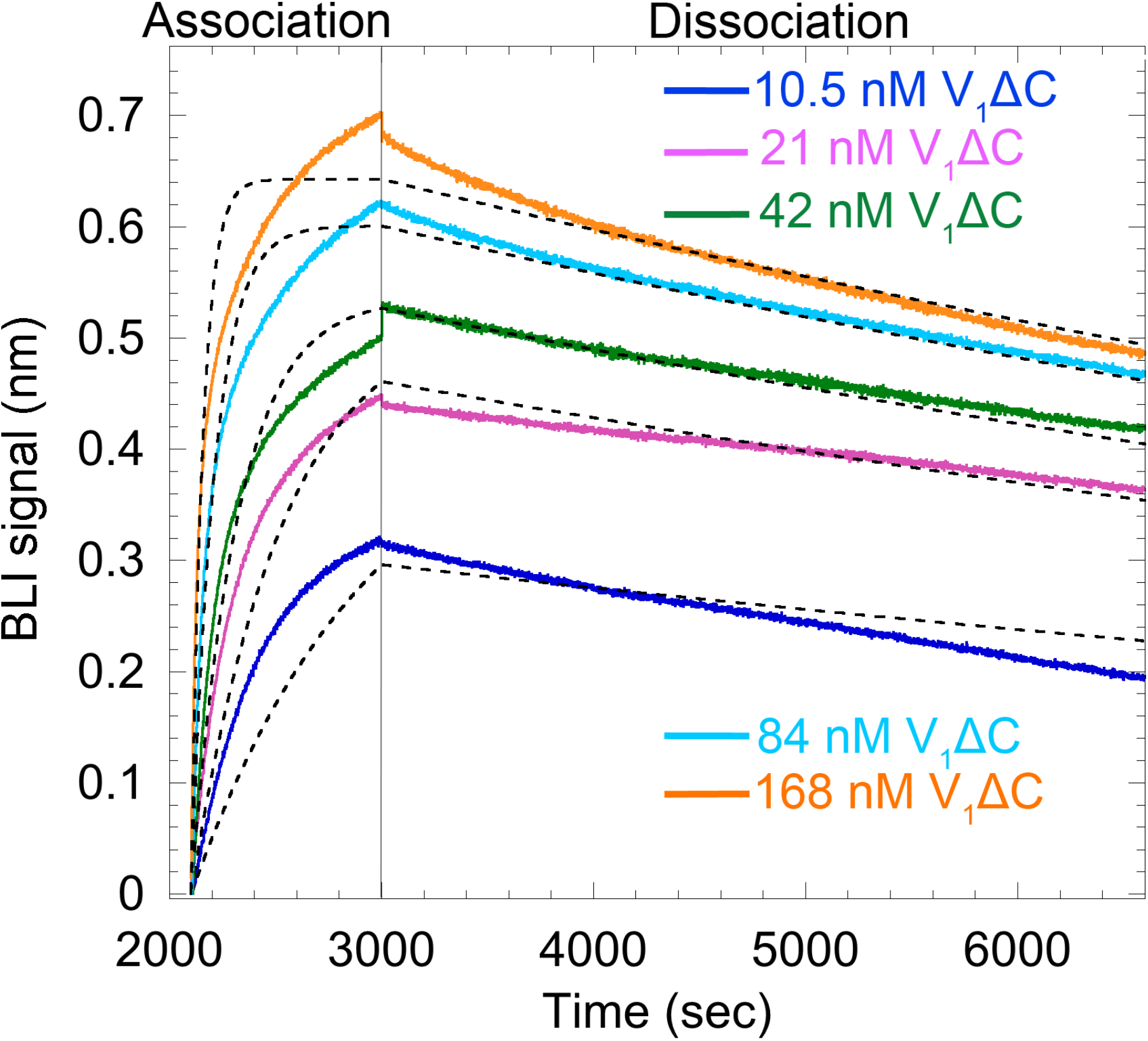
Binding of the C subunit to V_1_ΔC as analyzed by BLI. Anti-MBP antibody at 1 μg/ml was immobilized on anti-mouse IgG Fc capture (AMC) sensors. Anti-MBP containing sensors were dipped in 5 μg/ml MBP-C containing wells followed by wells containing 10.5, 21, 42, 84, and 168 nM of V_1_ΔC and buffer as control. All sensors were then dipped in buffer for dissociation. MBP-C dissociated from anti-MBP at a slow rate which was subtracted for the analysis of kinetic data. The association and dissociation curves were fit to a global single site model, giving a Kd of ~0.7 nM. A representative from two separate experiments is shown.

